# Single Cell Expression Data Reveal Human Genes that Escape X-Chromosome Inactivation

**DOI:** 10.1101/079830

**Authors:** Kerem Wainer-Katsir, Michal Linial

**Affiliations:** Department of Biological Chemistry, The Institute of Life Sciences, The Hebrew University of Jerusalem, ISRAEL

**Keywords:** X-inactivation, Allelic bias, RNA-seq, Escapees, single-cell

## Abstract

Sex chromosomes pose an inherent genetic imbalance between genders. In mammals, one of the female’s X-chromosomes undergoes inactivation (Xi). Indirect measurements estimate that about 20% of Xi genes completely or partially escape inactivation. The identity of these escapee genes and their propensity to escape inactivation remain unsolved. A direct method for identifying escapees was applied by quantifying differential allelic expression from single cells. RNA-Seq fragments were assigned to informative SNPs which were labeled by the appropriate parental haplotype. This method was applied for measuring allelic specific expression from Chromosome-X (ChrX) and an autosomal chromosome as a control. We applied the protocol for measuring biallelic expression from ChrX to 104 primary fibroblasts. Out of 215 genes that were considered, only 13 genes (6%) were associated with biallelic expression. The sensitivity of escapees' identification was increased by combining SNP mapping for parental diploid genomes together with RNA-Seq from clonal single cells (25 lymphoblasts). Using complementary protocols, referred to as strict and relaxed, we confidently identified 25 and 31escapee genes, respectively. When pooled versions of 30 and 100 cells were used, <50% of these genes were revealed. We assessed the generality of our protocols in view of an escapee catalog compiled from indirect methods. The overlap between the escapee catalog and the genes’ list from this study is statistically significant (P-value of E-07). We conclude that single cells’ expression data are instrumental for studying X-inactivation with an improved sensitivity. Finally, our results support the emerging notion of the non-deterministic nature of genes that escape X-chromosome inactivation.

## INTRODUCTION

Sex chromosomes pose an inherent genetic imbalance of expression between the genders (Lyon 1999; Graves 2006; Livernois *et al.* 2012). In order to ensure balanced expression, in mammals, one of the female’s X-chromosomes (ChrX) undergoes random inactivation (Chow and Heard 2009; Gabory *et al.* 2013). The random choice for an inactivated X-chromosome (Xi) (*i.e.,* the paternal or the maternal one) is completed at a very early phase of embryonic development (Dupont and Gribnau 2013; Petropoulos *et al.* 2016). A recent study on embryonic human cells revealed the dynamics of gene silencing throughout the first steps of embryology unti implementation (Petropoulos *et al.* 2016). Importantly, once this decision is made, the selected inactivated chromosome is deterministically defined for all descendant cells, and this choice is maintained throughout the organism’s life. This highly regulated process has been studied extensively (Avner and Heard 2001; Csankovszki *et al.* 2001; Wutz and Gribnau 2007; Briggs and Reijo Pera 2014).

Silencing and inactivation of Xi are maintained through epigenetic factors that drive the chromosome to possess a heterochromatin pattern (Balaton and Brown 2016). Initial silencing of X-chromosome is governed mainly by *XIST* (X-inactive specific transcript) (Penny *et al.* 1996), a non-coding RNA unique to placental mammals. *XIST* is a master regulator located at the X-inactivation center (XIC) (Pontier and Gribnau 2011). The gene is transcribed from Xi, and its RNA product acts in cis by coating the chromosome within a restricted chromosomal territory (Plath *et al.* 2002; Agrelo *et al.* 2009). *XIST* is also crucial for recruiting chromatin remodeling complexes (Bailey *et al.* 2000; Wutz *et al.* 2002; Gimelbrant *et al.* 2007; Vallot *et al.* 2013; Moindrot and Brockdorff 2016), resulting in an irreversible heterochromatinization (Brown and Robinson 2000; Augui *et al.* 2011; Barakat and Gribnau 2012). The epigenetic marks on the Xi include hypoacetylation and hypermethylation (e.g., H3K27me3) of promoter regions (Avner and Heard 2001; Balaton and Brown 2016). Additionally, the active X-chromosome (Xa) and Xi differ in their 3D structure (Marks *et al.* 2015). Apparently, chromosomal features such as loop boundaries and topologically associated domains (TADs) are important attributes in the dynamic of ChrX silencing process (Nora *et al.* 2012; Peeters *et al.* 2014; Deng *et al.* 2015).

Silencing does not apply to all genes in the inactivated X-chromosome. Interesting exceptions are genes that are shared between the sex chromosomes. These genes are located in regions, called pseudoautosomal regions (PARs) which are essential for a proper segregation of chromosomes during meiosis in males. In humans, PARs include 29 genes located at the tips of the X-chromosome, and are expressed from both alleles, similar to any autosomal chromosomes. In addition, other genes from Xi, called escapees, have the tendency to escape inactivation (Berletch *et al.* 2011). However, a substantial heterogeneity in the identity of these genes was reported among cells and experimental conditions (Carrel and Willard 1999). Escapees are mostly associated with evolutionarily young segments, presumably within the segment of ChrX that recently (on an evolutionary time scale) diverged from the Y-chromosome (Carrel and Willard 2005; Ross *et al.* 2005). The estimated fraction of escapees in human accounts for 15-20% of genes on ChrX (Disteche 1995; Balaton *et al.* 2015). Interestingly, many mouse homologous for human’s escapees are located in autosomal chromosomes (Berletch *et al.* 2015). Overall, the fraction of escapees in mouse is substantially smaller with respect to humans (Berletch et al., 2010).

Escaper genes in humans were mostly identified by indirect technologies (Peeters *et al.* 2014). In most instances, RNA expression levels in tissues were compared for males and females (Talebizadeh *et al.* 2006; Yasukochi *et al.* 2010). In other settings, differential RNA expression was measured from females with skewed X-chromosome inactivation (Cotton *et al.* 2013). Other methods focus on a cellular perspective including comparing healthy cells (46,X,X) to cells extracted from females with excess copies of ChrX (Sudbrak *et al.* 2001). An extensive catalog of escapee candidates was reported from mouse-human cell hybrids (Brown *et al.* 1997; Carrel and Willard 2005; Balaton *et al.* 2015). An additional approach for escapees’ detection considers the lack of methylation in CpG islands on Xi (Hellman and Chess 2007; Weber *et al.* 2007). In accord with an epigenetic view, a high-resolution mapping that compares the pattern of methylation in females with normal (45,X,X) and Turner (45,X) karyotypes was presented. The results substantiated the correlation between escapees and methylation pattern (Sharp *et al.* 2011). The varying expression levels of the candidate escapees may explain variations in phenotypes and clinical outcomes in women and men with an altered appearance of sex chromosomes (Lyon 2002). The ability for assigning specific alleles from Xa and Xi enables quantifying the statistical biases underlying imbalanced allelic expression. Furthermore, it was assumed that genes escaping X-inactivation have characteristic features for the absolute level of their expression (Zhang *et al.* 2013; Balaton and Brown 2016).

In this study, we present a protocol for RNA-Seq data that is specifically designed for single-cells. We focus on cells that can be distinguished by allelic SNPs via the information extracted from a reference genome. Based on the allelic expression of genes on X-chromosome combined with detailed information on parental chromosomes, we identify escapees and inactivated genes. We present general principles on the identification of escapees in view of cell types and diverse biological contexts. We also discuss the advantage and limitations of single cell genomics and transcriptomics to quantify allelic imbalance phenomena.

## MATERIALS AND METHODS

### Reference genome for the single cell primary fibroblasts

DNA-seq from female newborn primary fibroblast culture derived from umbilical cord tissue from newborns of western European origin was used (called UCF_1014). Data was extracted from EGAS00001001009 (https://www.ebi.ac.uk/ega/studies/EGAS00001001009) (Dimas *et al.* 2009; Borel *et al.* 2015). DNA was isolated and libraries were prepared by TruSeq DNA Kit (Illumina) and sequenced on two lanes of HiSeq2000 machine as 100 bp paired end reads. The DNA-seq data we extracted was realigned by BWA (Li *et al.* 2009) to the hg19 reference genome. Variation was called using GATK best practices procedure (Van der Auwera *et al.* 2013). For increasing the confidence of the analysis, the 2 VCFs were represented by one VCF using BCFtools utilities (Li 2011). In order to consider a SNP for further analysis, we required that a SNP to appear in both VCFs (bcftools isec -n+2 -o UCF_1014.vcf -O v-p UCF_1014/ -w1). Only heterozygous variations were compiled for further analysis.

### RNA-Seq data for the single cell fibroblasts

RNA-seq of single cells were obtained from EGAS00001001009 as above (Borel *et al.* 2015). As described in (Borel *et al.* 2015), single cells were harvested and cDNA was prepared in the C1 Single-Cell Auto Prep system (Fluidigm). The preparation of RNA included pre-amplification with Advantage-2 PCR Kit. Libraries were made via Nextera XT DNA Kit (Illumina) and sequenced on HiSeq2000 machine as 100 bp paired end reads. For consistency, we choose to analysis only cells that were amplified by an identical protocol (i.e., 22 cycles of PCR, total 104 cells). The RNA-seq reads were cleaned using Trimmnomatic (Bolger *et al.* 2014). RNA-seq was realigned to UCSC hg19 reference using TopHat2, a splice junction mapper for RNA-Seq reads (Langmead and Salzberg 2012), allowing 2 mismatches with no gaps. Repeated reads were marked using Picard, and RNA-seq was indexed using SAMtools.

### Allelic imbalance analysis in the single cell fibroblasts

All reads from each BAM alignment file were counted against the SNPs on the VCF using Allelcounter-master (Castel *et al.* 2015). Reference and Alternative assigned reads were counted. Results for chromosome-X and chromosome-17 (ChrX, Chr17) were further analyzed by R. A cell specific threshold for a minimal expression level was set. A threshold of 0.00002% of the aligned reads in a BAM file was used, which on average accounts for ~5 reads per SNP. SNPs that were mapped with a lower number of reads were not included in the analysis. Allelic ratio (AR) was calculated for each of the informative SNPs. AR is defined as the ratio based on the number of reads matched to the alternative SNP (#Alt) divided to the sum of the reads for both alleles, the reference (Ref) and Alt (#Ref + #Alt). As the origin of each allele is unknown in the case of the non-phased genome, only genes with evidence for biallelic expression from the same cell on ChrX, and are supported by multiple evidence are considered escapees.

### Reference genome for the single cell lymphoblasts

The reference genome used for GM12878 cell line is the diploid NA12878 genome (version Dec 16, 2012, from http://alleleseq.gersteinlab.org/). This genome of the GM12878 cell line is based on hg19, with 4,330,326 SNPs and 829,454 INDELs. The variant list is based on HiSeq 64x sequencing call set from the BROAD institute. Details are available in ftp://gsapubftp-anonymous@ftp.broadinstitute.org/bundle (Genomes Project *et al.* 2010; Mills *et al.* 2011). The diploid genome was extracted from http://sv.gersteinlab.org/NA12878_diploid/NA12878_diploid_genome_2012_dec16.zip.

The allelic specific assignment was based on the computational pipeline AllelSeq (Rozowsky *et al.* 2011) and findings from ChrX and Chr17 are reported. The sequences for these chromosomes, each with paternal and maternal versions were used as a default, unless otherwise mentioned. Data were collected into one FASTA file that was indexed by Bowtie2 (Langmead *et al.* 2009).

### Mapping allelic specific SNPs for the single cell lymphoblasts

In addition to the diploid genome, we used a VCF file containing all known SNPs for the selected cell line. The file is available in http://sv.gersteinlab.org/NA12878_diploid/CEUTrio.HiSeq.WGS.b37.bestPractices.phased.hg19.vcf.gz. From the VCF file heterozygous SNPs having two haplotypes on ChrX and Chr17 were extracted. All SNPs were assigned to the canonical transcripts according to the compiled list available from the UCSC known gene list (Raney *et al.* 2014).

Remapping of the VCF coordinates with those of the diploid NA12878 genome (version Dec 16, 2012) was done using a mapping protocol for assigning positions on paternal and maternal haplotypes. The procedure uses Pearson's FASTA36 program (from http://faculty.virginia.edu/wrpearson/fasta) and local alignment BLAST extended for 500 nucleotides at each side of a SNP, for each haplotype. Activating LocalAlign function from Matlab Bioinformatics Toolbox (www.mathworks.com/products/bioinfo/) was used for further verification of the SNPs’ coordinates. For comparing sequences, a window of 100 nucleotides centered at the candidate SNP was created.

SNPs were verified for having a unique mapping on the genome. A strict mapping was based on matching the sequence into a window of 201 nucleotides for the paternal and maternal SNP alleles. In each of these segments, the SNP allele occupied the indexed nucleotide 101. From all 201 nucleotides long sequences, we created a FASTA file that was aligned with 'no gaps' to the full genome using Bowtie2. Only SNPs uniquely aligned to the genome (no 'XS:i' flag) were included in the analysis. After the verification step, we end up collecting 12,856 and 14,244 SNPs that are successfully mapped to ChrX and Chr17, respectively. These SNPs are represented in our SNP list.

Chromosomal locations of genes were obtained from UCSC hg19 GTF file provided by TopHat2. Converting chromosomal locations to parental chromosomal locations was performed using LiftOver available at UCSC toolbox (Rosenbloom *et al.* 2015). Paternal and maternal chain files for the process were downloaded along with the complete genome. The conversion created a GTF file with ChrX and Chr17 maternal and paternal locations of genes. This GTF was then used for TopHat2 alignment. The final step in the mapping includes creating a reference GTF file for the positions of the SNPs list on each chromosome (ChrX and Chr17). This reference GTF file contains all SNP locations (called GTF_SNP). Formally, in the GTF file, each SNP was considered as having a match with either a paternal feature on paternal chromosome or a maternal feature on maternal chromosome (Castel *et al.* 2015). RNA-Seq alignments to the entire genome (instead of to ChrX and Chr17) was performed for a represented single cell (SRR764802, see Supplemental Table S4). The results were practically identical for the two alignment schemes.

### RNA-Seq data for the single cell lymphoblasts

RNA-Seq experiments from GM12878 lymphoblastoid cell-line single cells were used as the source for allelic assignment (Marinov *et al.* 2014). GM12878 cells were originated from female’s blood of a European ancestry. The cells have a normal karyotype and sequencing was performed using Illumina HiSeq 2000 with 100-mer reads. Libraries were constructed by SMART-Seq protocol (Ramskold *et al.* 2012). Data from 25 single cells RNA-Seq files were downloaded from http://www.ncbi.nlm.nih.gov/geo/query/acc.cgi?acc=GSE44618. Additional data include pool of 30 and 100 individual cells’ paired end RNA-Seq data files. The same pipeline was used for the pool and single cells. FASTQ files were cleaned using FASTX (http://hannonlab.cshl.edu/fastx_toolkit/index.html). We have used FASTA_Clipper protocol with rigorous parameters for trimming out low-score positions (fastq_quality_trimmer -Q33 -t 25 -l 25 –i). SMART adaptors were removed from the sequenced fragments (-Q33 -a AAGCAGTGGTATCAACGCAGAGTACTTTTTTTTT -l 25 -i). Each trimmed file was aligned to the paternal and maternal chromosomes using TopHat2 (Kim *et al.* 2013) with the following parameters: no mismatches '-N 0', no spaces '--read-gap-length 0 ', and a sensitive alignment addition '--b2-very-sensitive' that checks every sequence several times to improve sensitivity and accuracy. We eliminated non-unique mapping (using ‘NH:i:1’ flag), with thresholds for alignment length >= 50 nucleotides.

HTSeq pipeline (Anders *et al.* 2015) was used for read counting. We used GTF_SNP as a reference for the HTSeq positions of interest in the genome. HTSeq counted how many of each of the features overlapped the SNPs locations indicated in the GTF_SNP file. We included identified SNPs from the same fragment that were referred by HTSeq as “ambiguous” (applies for rare instances of closely positioned SNPs). Overall, HTSeq listed for each SNP, the location and number of perfectly aligned reads to maternal or paternal alleles. We activated SamTools for viewing the aligned reads (Li *et al.* 2009).

### Allelic imbalance analysis for the single cell lymphoblasts

The single cells female lymphoblasts are clonal and therefore, inactivation is associated with a particular X-chromosome which is shared by all the cells. An expression ratio was calculated for each SNP as the number of parental reads over the total reads at this position (i.e., reads from maternal and paternal genomes). As cells differ in the number of successfully mapped reads, a cell-specific threshold was applied. We only consider SNPs with more than 0.001% of mapped reads from the sum of ChrX and Chr17 unique alignments. This threshold accounts for 4-20 reads/cells, with an average of 7 reads per cell. Replacing the cell-specific threshold by a predetermined read value (e.g., >=7 reads) has only a minor effect on the results. Informative SNPs are the collection of SNPs having a heterologous position within canonical transcripts with expression level above the cell-specific threshold. We assigned 4 types of labels according to the expression ratio associated with each informative SNP: (i) paternal; (ii) biallelic expression leaning towards paternal; (iii) biallelic expression leaning towards maternal and (iv) maternal. The labels are set by quartiles. Specifically, SNPs with an allelic ratio of >0.75 and =<0.25 were labeled paternal and maternal, respectively. SNP is labelled “biallelic maternal” for 0.25<allelic ratio=<0.5, and as “biallelic paternal” for 0.5<allelic ratio<=0.75. For some analysis, we combined labels (ii) and (iii). These SNPs were labeled ‘balanced’ for expression ratio of 0.25<ratio<=0.75. Each informative SNPs is counted as a data-point (DP). An identical labeling thresholds were applied for single cells and for pool of cells (marked Pool 30 and Pool100 for 30 and 100 pool of cells, respectively).

Expression data from single cells are sparse and prone to noise. Therefore, we applied two complementary protocols. For the strict protocol, we consider genes that are supported by at least two DPs. Multiple DPs may represent informative SNPs from multiple cells or informative independent SNPs in a gene, from the same cell. Due to the sparseness of the informative SNPs, multiple DPs were mostly obtained from different cells. We defined a DP score to estimate the level of support for different genes and as a baseline for comparing information from multiple resources. The DP score is a simple summation of the DP labels. Where the SNP labeled as maternal is scored 0 and the paternal or balanced expression are scored 1. We consider escapees as genes with DP score >1. We represent the score as the fraction of DP score out of all DPs present.

For the relaxed mode, the identification between inhibited and escapee genes is done by counting the number of reads per informative SNPs. Each gene is associated with the sum of the reads overlapping the informative SNPs within its canonical transcript. A gene will be considered as an escapee if the number of paternal reads is >=6. For inhibited genes, we set higher threshold in which the allelic ratio is <0.05 with maternal reads >=100).

### Comparison to a unified annotated ChrX gene catalog

There are 1144 known genes in ChrX (Rosenbloom *et al.* 2015). These genes were annotated according to 9 phenotypes according to (Balaton *et al.* 2015). The labels are PAR, escapee, mostly escapee, variable escapee, mostly variable escapee, genes associated with strongly conflicting results, inhibited genes, mostly inhibited and genes having no data (Balaton *et al.* 2015). The annotations are based on a careful analysis according to major publications combining numerous indirect measurement for escapee and inhibited gene identification (Balaton *et al.* 2015). From all genes in ChrX, 45% have no data, 40% are inhibited-related, 4% have conflicting evidence and the rest carry escapee-related annotations. We consider escapee-related benchmark as all genes that carry escapee annotations, including the set with conflicting evidence and PAR genes (total 168 genes).

### Statistical analysis

Hypergeometric probability between our single cell results and the annotated catalog was calculated by comparing the correspondence of any two lists of escapees. We used standard notations of N, k, n and x: N symbolizes all genes that are expressed from ChrX with the subjected phenotypes as defined above (Balaton *et al.* 2015); n is the number of escapees we identified by any of our protocols; x the number of genes in our list that match the literature-based escapee list in k. P(x) is the probability that an n-trial will result in a value that is >= x.

## RESULTS AND DISCUSSION

### Pipeline for X-inactivation for single cells

We set out to identify escapee genes by analyzing RNA-Seq gene expression from single cells. Analyzing single cells presents a clear advantage over indirect methods since it ensures that inactivation will be associated with one specific chromosome (Xi) in each cell. We considered two cases for single cell’s data. For most instances, detailed data on SNP variations from parental genomes are unavailable. However, in the cases that a diploid parental genome is available, deep sequencing of cells’ transcriptome provides essential information for determining the parental origin of SNPs and other informative variants. Thus, enabling escapees’ identification.

Figure 1 illustrates the pipeline used for our analysis. In a nutshell, we analyzed a large collection of single-cell transcriptomes (104 cells, primary female human fibroblasts). For this collection, parental phased genomes were unavailable and therefore, escapees were determined by evidence for biallelic expression (Figure 1A). Figure 1B emphasizes quantitative differences between ChrX and Chr17. Specifically, Chr17 is richer in genes as compared to ChrX (14.3 and 5.3 coding genes per 1M nucleotides, respectively). Additionally, we analyzed a collection of RNA-Seq data for lymphoid cell-line (25 clonal lymphoblast cells), for which parental genomes are known (Figure 1C). Identifying escapees relies on two complementary protocols. Under a strict protocol, each informative SNP is labeled as a Data Point (DP) by its allelic preference (see Materials and Methods). In the relaxed protocol, an escapee gene is defined according to the sum of the mapped reads that cover informative SNPs and are expressed from the Xi chromosome (Figure 1C).

**Figure 1.**
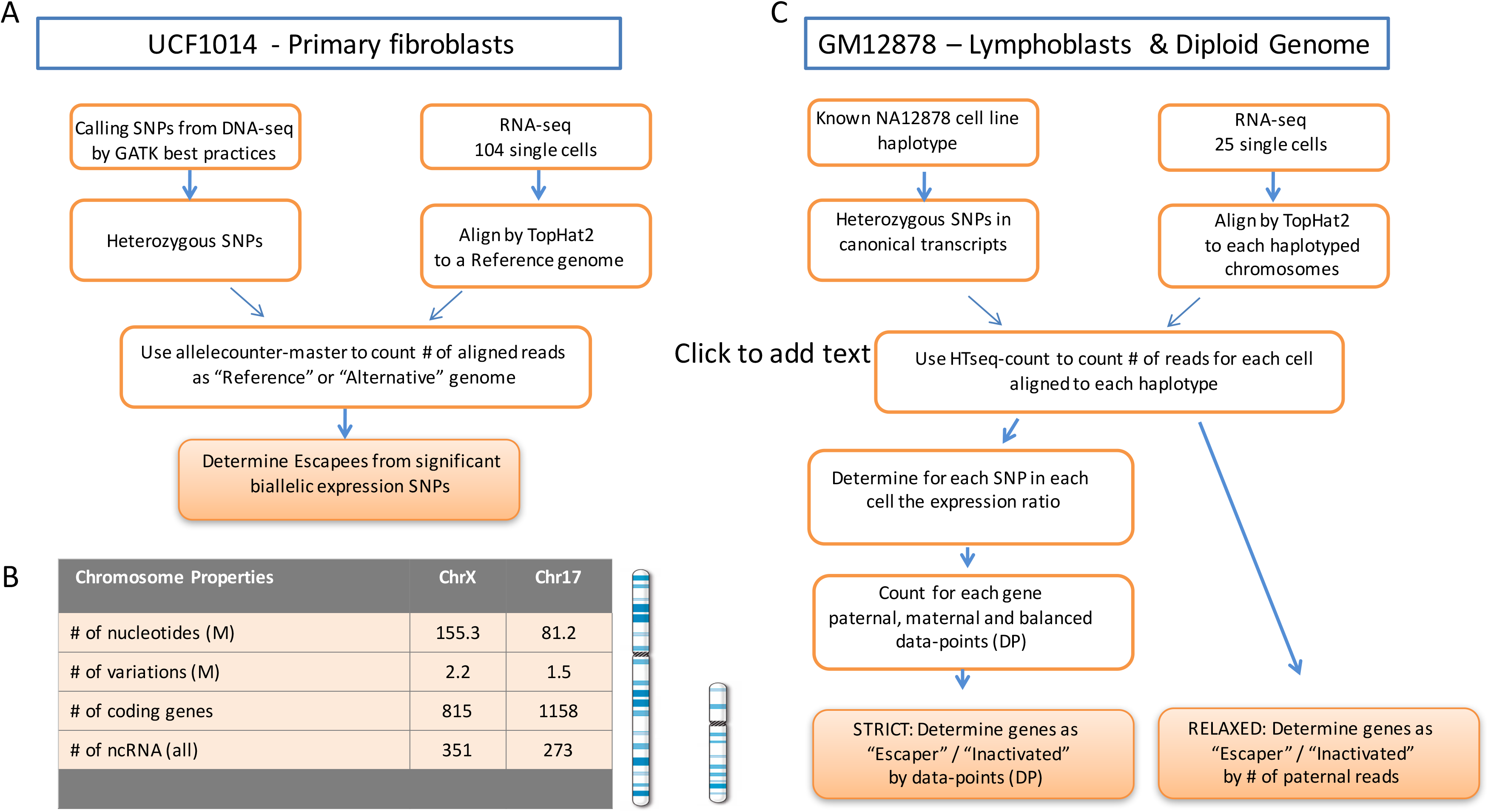
A workflow for identifying escapee genes from single cells RNA-Seq data. (A) The protocol applied for identifying escapees using DNA-seq from 104 single cell RNA-seq data. Identical protocols are applied to ChrX and Chr17. (B) Quantitative properties summary for ChrX and Chr17 in view of the number of coding, non-coding genes, and the physical properties of the ChrX and Chr17. The chromosomes schematics are shown. (C) An outline scheme describing strict and relaxed protocols that were used for analyzing 25 single cell lymphoblasts with known haplotypes.

### Single cell biallelic expression in primary human fibroblasts

Large scale transcriptomic data from female human fibroblasts were used to assess X-Chromosome Inactivation (XCI), and the phenomenon of genes that escape inactivation (Borel *et al.* 2015). The reliability of sequence data from individual cells was extensively studied and will not be further discussed (Marinov *et al.* 2014). A total of 104 high-quality data from single cells were analyzed (for sequencing depth and mapping results, see Supplemental Table S1). For each cell, informative, heterologous SNPs are listed, and each of these SNPs was assigned with a label according to the expression ratio for the two alleles (see Materials and Methods, Supplemental Table S2). The number of informative SNPs on Chr17 and ChrX in individual cells are correlated (r = 0.62, p-value = 2.78E-12) supporting the accuracy of the mapping protocol (Supplemental Figure S1).

Figure 2 summarizes the findings from Chr17 and ChrX according to allelic ratio (AR) from single-cells primary fibroblasts. Expression ratio of 0 associated with an exclusive expression from the alleles represented by the reference genome (Figure 2). The read-mapping is somewhat biased towards the reference genome, as previously reported (Degner *et al.* 2009; Panousis *et al.* 2014). As expected, Chr17 and ChrX display completely different proportion for biallelic gene expression. While biallelic expression accounts for 24% of the SNPs’ occurrence (6100/25,324) for Chr17, it accounts for <9% in ChrX (870/9795, Figure 2A-2B). When only biallelic expression is considered for both chromosomes (Figure 2C-2D), only Chr17 displays a distribution that matches a balanced appearance (centered around x-axis=0.5).

**Figure 2.**
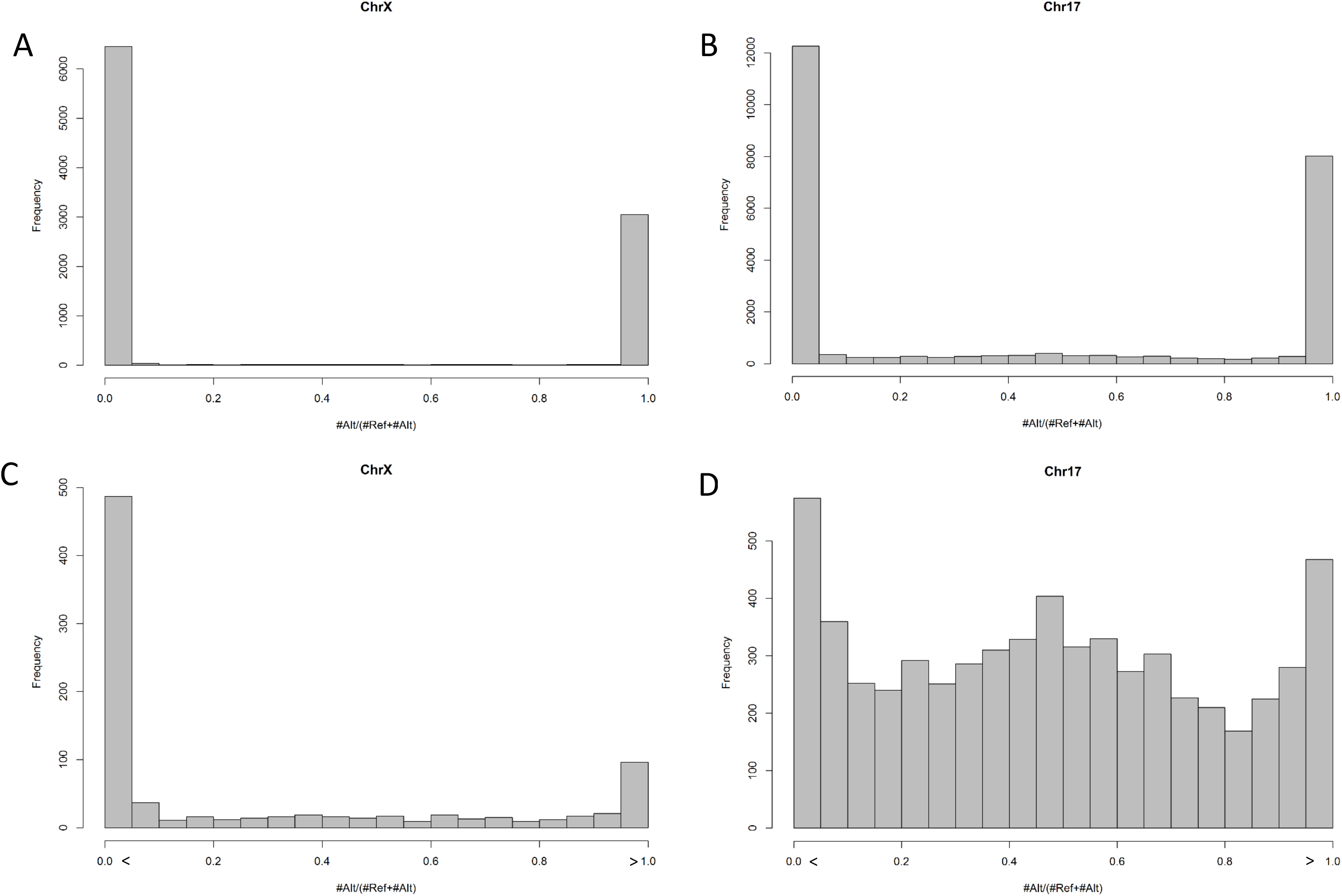
The distribution of the allelic ratio (AR) for each SNP as a fraction of the assignments for Alternative (Alt) or Reference (Ref) alleles. AR considers the fraction of each allele in view of total counts associated with this SNP. X-axis ranges from 0 to 1.0, where 0 indicates that all assignments that are associated with the Ref allele. The distributions of the allelic ratio for ChrX (A) and Chr17 (B) are shown. As most of the SNPs are assigned as 0 or 1, we zoomed on the informative SNPs that are expressed from both alleles (i.e., 0 < AR <1). The zoomed analysis for AR distribution for ChrX (C) and Chr17 (D) is shown. Note that there is a substantial number of informative SNPs with a mixed expression for autosomal Chr17 while for ChrX it is a rare phenomenon.

Table 1 summarizes the list of escapees derived from the primary single fibroblasts. Only 13 genes are marked as escapee candidates. These genes are characterized by a significant “balanced expression” signal. A maximal support is linked to *ZFX* (Zinc finger X-chromosomal protein) and *SMC1A* (Structural maintenance of chromosomes protein 1A). The number of genes that exhibit biallelic expression in Chr17 is 10 fold higher (142 genes). The mark difference in the abundance of biallelic expression from ChrX and Chr17 (Figure 2, Supplemental Table S3) is a strong indication of the stability of XCI phenomenon in primary isolated cells. Actually, without phasing, only genes that show genuine biallelic expression in the same cell can be securely identified as escapee candidates. Noticeably, PAR genes that are characterized by a biallelic expression were not identified. This is an outcome of our mapping protocol which was performed on a male genome. Obviously, sequences that are identical between ChrX and Y-chromosome, including the PAR genes were eliminated on the basis of not having a unique mapping (see Materials and Methods). The lack of parental chromosomes enforces alignments to a reference genome. Using a single reference genome, was shown to severely affect alignment results (Satya *et al.* 2012). We conclude that the primary origin of the analyzed fibroblasts and the lack of phased parental chromosomes reduced the discovery rate for escapees.

**Table 1.** List of escapees identified by the strict protocol from fibroblast single cells’ transcriptome.

| Gene symbol | ChrX Band | Comment | <sup>a</sup>Ratio (DPs) E-score | <sup>b</sup>Ratio (reads) E-Score | Name / Function |
| --- | --- | --- | --- | --- | --- |
| LAMP2 | Xq24 |  | 0.016 | 0.310 | Lysosomal-associated membrane protein 2 |
| ZRSR2 | Xp22.2 |  | 0.071 | 0.332 | Zinc finger (CCCH type), RNA-binding motif and serine/arginine rich 2 |
| TCEAL4 | Xq22.2 |  | 0.056 | 0.335 | Transcription elongation factor A (SII)-like 4 |
| HDHD1 | Xp22.2 |  | 0.039 | 0.346 | Haloacid dehalogenase-like hydrolase domain containing 1 |
| XIAP | Xq25 |  | 0.025 | 0.398 | X-Linked Inhibitor of Apoptosis, E3 Ub Ligase |
| C1GALT1C1 | Xq24 |  | 0.020 | 0.475 | C1GALT1-specific chaperone 1 |
| ZFX | Xp22.11 |  | 0.186 | 0.528 | Zinc Finger Protein, X-Linked |
| SMC1A | Xp11.22 |  | 0.236 | 0.541 | Structural Maintenance of Chromosomes 1A |
| DDX3X | Xp11.4 |  | 0.052 | 0.546 | DEAD (Asp-Glu-Ala-Asp) Box Helicase 3, X-Linked |
| RBMX | Xq26.3 |  | 0.042 | 0.613 | RNA binding motif protein, X-linked |
| LOC550643 | Xp11.21 | ncRNA | 0.030 | 0.624 | Uncharacterized LOC550643 |
| RBM3 | Xp11.23 |  | 0.035 | 0.694 | RNA binding motif (RNP1, RRM) protein 3 |
| EDA2R | Xq12 |  | 0.056 | 0.777 | Ectodysplasin A2 receptor |

<sup>a</sup>Escaper Score according to the strict protocol (DPs) for 104 single fibroblast cells. The score (0-1.0) is based on scoring genes by balanced data points (DPs) out of total DPs. <sup>b</sup>Escaper Score according to the relaxed (reads' count) protocol. The ratio (0-1.0) is based on the fraction of the Alternative allele in view of the total sum of reads per gene.

### Biallelic expression in clonal human lymphoblasts

We set to increase the information that can be extracted from single cells by focusing on clonal cell lines. To this end, we analyzed female RNA-Seq from 25 single lymphoblast cells (clonal, GM12878, Supplemental Table S4). The activated X-chromosome (Xa) of GM12878 cells is associated with maternal origin (Marinov *et al.* 2014). We benefited from the availability of a diploid genome with paternal and maternal reference chromosomes (NA12878, see Materials and Methods). The clonal nature of these 25 cells allows overcoming the unavoidable cell-cell variability without being masked by the stochastic nature of XCI (in contrast to primary fibroblasts, Figure 2). Similar to the observation shown for primary fibroblasts (Supplemental Figure S1), a strong correlation between the number of informative SNPs in Chr17 and ChrX in individual cells was observed (is r = 0.95, p-value = 5.99E-13, Supplemental Figure S2). Note that about a quarter of the cells poorly contribute to the analysis and carry <10 informative SNPs per cell for ChrX. The most informative 20% of the cells contribute >30 informative SNPs each (Supplemental Table S5).

Figures 3A-3B show the partition of data points (DPs) assigned for ChrX and Chr17 for each of the cells according to the expressed allele (maternal, paternal or ‘balanced expressed’, Supplemental Table S5). In all cells, the maternal expression from the active ChrX dominates. Accordingly, the paternal chromosome represents Xi in all the analyzed clonal cells. It is also evident that most cells, excluding a few low expressers, include a substantial fraction of non-maternal alleles, thus escaping from X-inactivation is exposed as a strong phenomenon at cell level. In contrast, Chr17 of single cells shows an equal contribution of both alleles with a high fraction of biallelic expression (Figure 3B).

**Figure 3.**
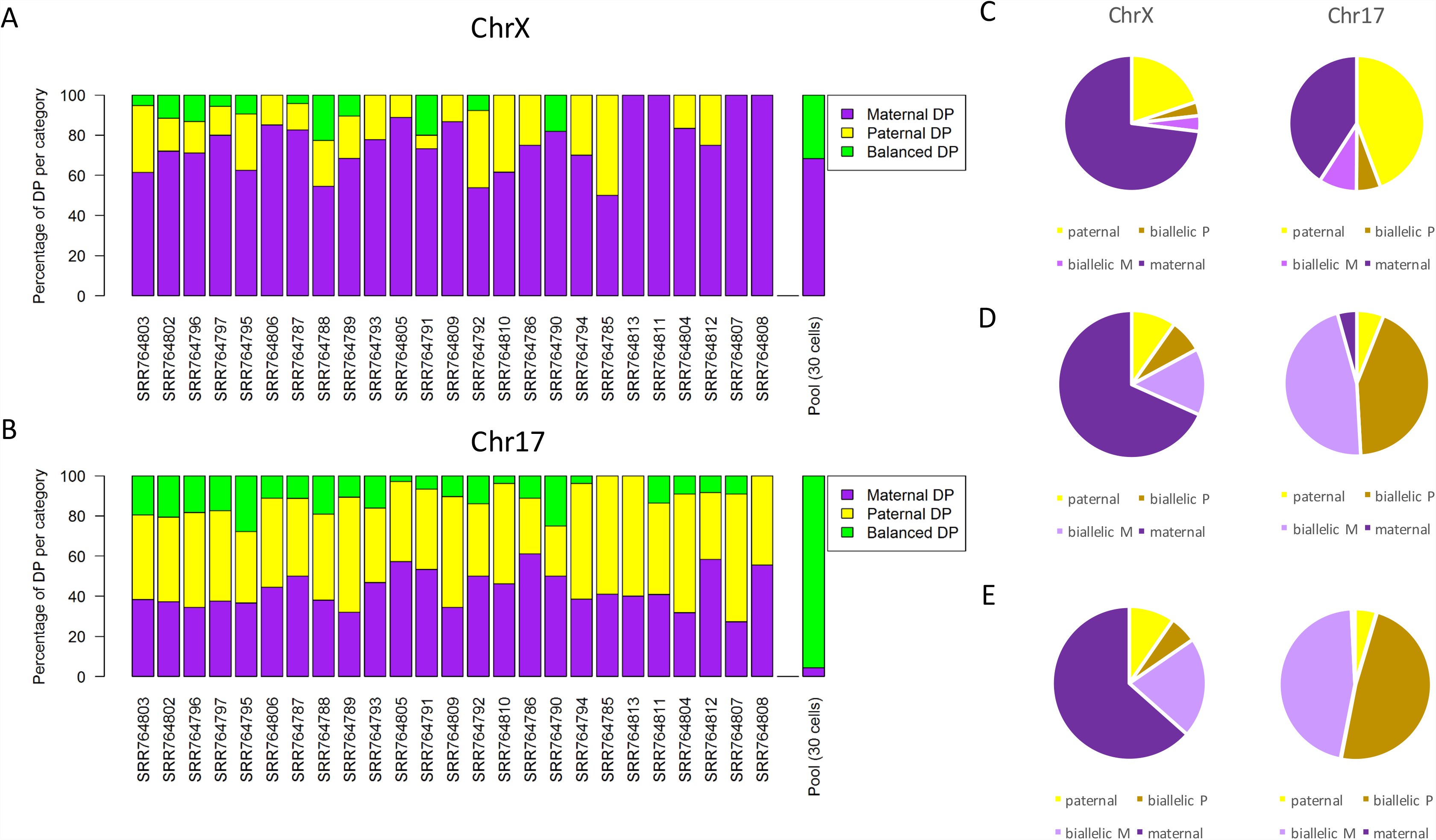
Quantifying the labels of informative SNPs from 25 single cell lymphoblasts. The sequencing depth for each of the analyzed cells and pool data are listed in Supplemental Table S4. Each cell is partitioned according to its categorical DPs on ChrX (see Materials and Methods). (A) The partition of DP labels for 25 single cells and Pool30 for ChrX is shown. The maternal, paternal and balanced expression are colored purple, yellow and green, respectively. (B) The partition of DP labels for 25 single cells and Pool30 for ChrX is shown. Color code for the expression labels is as in (A). A summary of the partition of labels for all 25 single cells on ChrX and Chr17 (C), pool30 (D) and pool100 (E) are shown. Each quantile of the AR values is differently colored. The data is based on 232 informative SNPs for ChrX and 455 informative SNPs for Chr17. The Pool30 data consists of 41 SNPs on ChrX and 116 on Chr17. The Pool100 consists of 52 SNPs on ChrX and 130 SNPs on Chr17.

The advantage of identifying escapees from single cells’ RNA-seq was tested with respect to data derived from pool of cells (Figures 3A-3B pool, Supplemental Table S6). We analyzed pools composed of 30 and 100 individual cells (Pool30 and Pool100 respectively). Specifically, the number of mapped reads for Pool30 is only 12% (5512 vs. 45841 reads) of the unified number of reads derived from all 25 single cells. A similar trend is apparent in Pool 100. In view of the informative SNPs, we collected evidence for 41 (Pool30) and 52 (Pool100) informative SNPs as compared to 235 labeled SNPs from individual cells. When the same analysis was applied to Chr17, the fraction of ‘balanced expression’ was substantially higher (compare Supplemental Table S5 and Supplemental Table S6). We conclude that the pool data provides a reduced sensitivity and a limited discovery rate with respect to the single cells data. This observation is not a result of the depth of the RNA-Seq (Supplemental Table S4).

Figures 3C-3E unify individual cells and present DP-centric results. It shows a partition of DPs for 25 single cells (Figure 3C) Pool30 (Figure 3D) and Pool100 (Figure 3E) for ChrX and Chr17. Importantly, the almost equal appearance of maternal and paternal DPs is shown for Chr17 (50%, 45% and 53% for 25 single cells, Pool30 and Pool100, respectively). These results are expected for gene expression from any autosomal chromosome. In contrast, most SNPs of ChrX are labeled maternal, in agreement with the origin of Xa in GM12878 lymphoid cells. Evidence for paternal expression from single cells accounts for 23% (Figure 3D, based on 232 informative SNPs). For the pool analysis, the paternal evidence accounts for only 15% and 16% of the DPs Pool30 and Pool100, respectively.

Importantly, data from an autosomal chromosome (Chr17) from single cells exhibit a strong tendency for mono-allelic expression. This observation reflects the phenomenon known as “transcriptional bursting” (Dar *et al.* 2012; Borel *et al.* 2015). An allele-specific expression burst prevails in single cells low-expressing genes. Pool data from autosomal Chr17 show an increase in the fraction of “balanced expressed” from 15% to 94% (compare single cells to Pool100). In contrast, the phenomena of inactivation of the X-chromosome (XCI) is reflected by a substantially reduced fraction of SNPs with “balanced expression” (18-27%, Figure 3C-3E).

### Expression of escapee genes from lymphoblasts

DPs in each single cell (Supplemental Table S5) and for pool data (Supplemental Table S6) were labeled according to the parental expression. We combined the evidence for a gene by unifying the DPs for a gene into an Escaper Score (Supplemental Table S7 and Supplemental Table S8). Figure 4A is a gene-centric view for ChrX by the number of DPs per gene. For improving the reliability of the identification, we requested multiple DPs as support. Activating the strict analysis protocol (as in Figure 1C) results in 64 genes on ChrX (25 escapees and 39 inhibited genes). For many of these genes, the number of DPs that support the genes' identity is rather small (Supplemental Table S7). Exceptions are *ZFX*, *CD99*, and *SLC25A6*, which are supported by 24, 25 and 36 DPs, respectively. Assuming that Chr17 has no significant allelic biases, out of 262 informative genes, only 16 genes are uniquely labeled paternal, and another 18 are labeled maternal (Figure 4B). Not surprisingly, these genes have only a few supporting DPs. Based on these observations, one can estimate false positives as 6% and negatives rates as 7% for ChrX escapers’ assignments.

**Figure 4.**
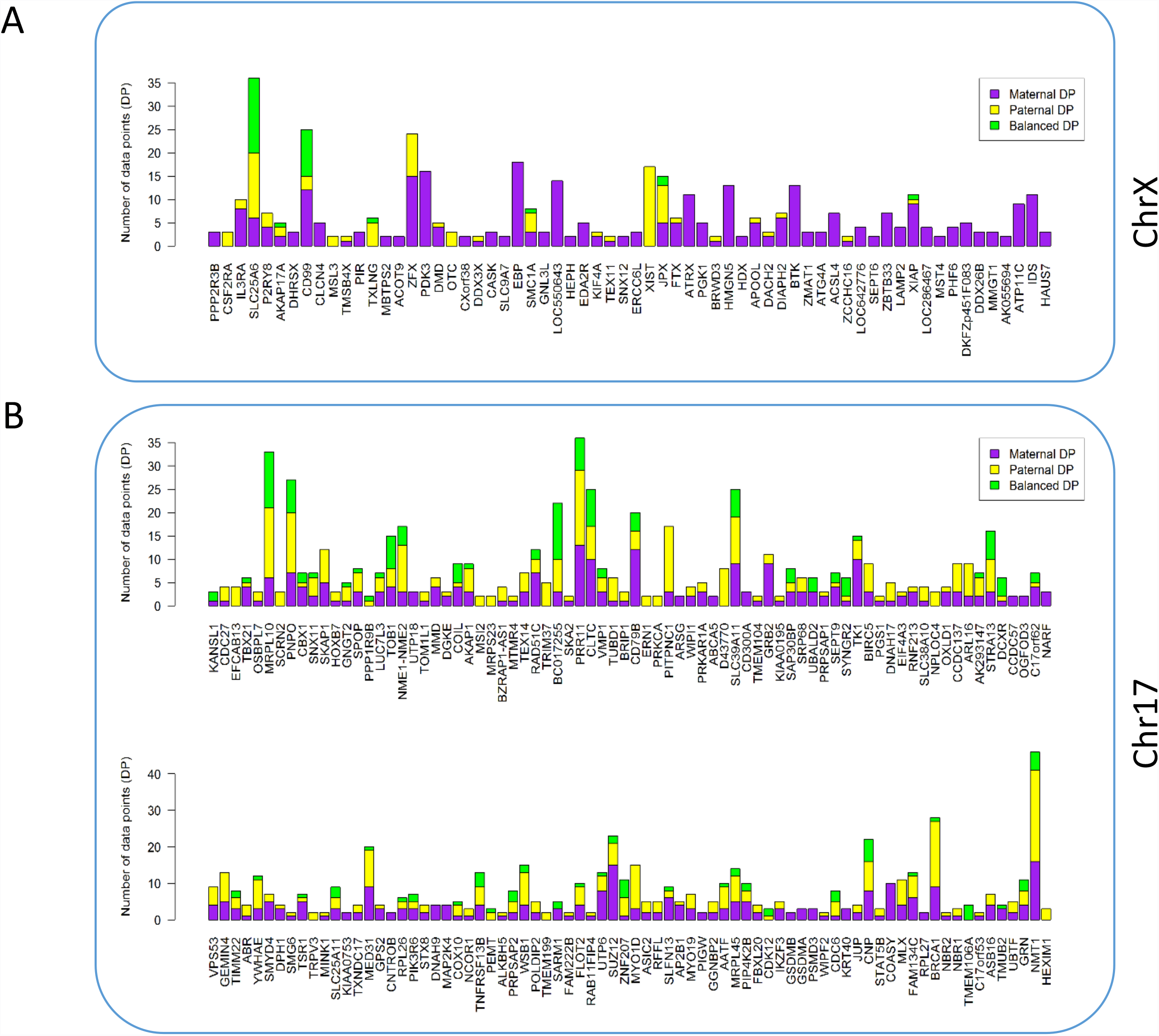
A gene-centric partition of DPs from 25 single lymphoblast cells. The difference in the partition of parental SNPs for ChrX (A) and Chr17 (B) according to the number of DPs is shown. The listed genes are those that are supported by multiple DPs. The color code is according to the DP label as paternal, maternal and “balanced expressed”. The 58 genes in ChrX and 262 genes in Chr17 are listed according to the order on the respected chromosome.

Candidate escapees along with their Escapee Score are listed in Table 2. In order to differentiate genuine escapees from false assignment, we revisited the appearance of PAR genes among the identified escapers (Table 2). We confidently identified by the property of biallelic expression 6 out the 7 expressed PAR genes (85.7% accuracy). This high discovery rate is in agreement with our estimation for the false negative rate. To further test the reliability of the mapped reads for ChrX, we tested the coherence in DPs’ labels in genes that are supported by multiple informative SNPs in a single cell (47% of genes, Supplemental Table S5). The assumption is that for the same gene in an individual cells the DPs are expected to be consistently labelled. Indeed, for ChrX, we confirmed the consistent labels among all SNPs that were associated with all genes (with one exception, JPX).

**Table 2.** List of escapees identified by the strict and relaxed protocol from lymphoblast single cells’ transcriptome.

| Gene symbol | ChrX Band | Comment | <sup>a</sup>Ratio (DPs) E-score | <sup>b</sup>DPs support | <sup>c</sup>Ratio (reads) E-Score | Name / Function |
| --- | --- | --- | --- | --- | --- | --- |
| PLCXD1 | Xp22.33 | PAR | - | S | 1 | PI Specific Phospholipase C X Domain Containing 1 |
| CSF2RA | Xp22.33 | PAR | 1 |  | 1 | Colony Stimulating Factor 2 Receptor Alpha |
| IL3RA | Xp22.33 | PAR | 0.2 |  | 0.077 | Interleukin 3 Receptor, Alpha (Low Affinity) |
| SLC25A6 | Xp22.33 | PAR | 0.833 |  | 0.532 | Solute Carrier Family 25 Member 6 |
| P2RY8 | Xp22.33 | PAR | 0.429 |  | 0.495 | Purinergic Receptor P2Y. G-Protein Coupled |
| AKAP17A | Xp22.33 | PAR | 0.6 |  | 0.656 | A Kinase (PRKA) Anchor Protein 17A |
| CD99 | Xp22.33 | PAR | 0.52 |  | 0.376 | CD99 Antigen |
| MSL3 | Xp22.2 |  | 1 |  | 1 | Male-Specific Lethal 3 Homolog (Drosophila) |
| FRMPD4 | Xp22.2 |  | - | S | 1 | FERM And PDZ Domain Containing 4 |
| TMSB4X | Xp22.2 |  | 0.5 |  | 0.118 | Thymosin |
| SYAP1 | Xp22.2 |  | - | S | 1 | Synapse Associated Protein 1 |
| TXLNG | Xp22.2 |  | 1 |  | 0.865 | Taxilin Gamma |
| ZFX | Xp22.11 |  | 0.375 |  | 0.140 | Zinc Finger Protein, X-Linked |
| DMD | Xp21.1 |  | 0.2 |  | 0.259 | Dystrophin |
| OTC | Xp11.4 |  | 1 |  | 1 | Ornithine Carbamoyltransferase |
| DDX3X | Xp11.4 |  | 0.5 |  | 0.45 | DEAD (Asp-Glu-Ala-Asp) Box Helicase 3, X-Linked |
| KDM6A | Xp11.3 |  | - | S | 0.995 | Lysine Demethylase 6A |
| KDM5C | Xp11.22 |  | - | S | 1 | Lysine Demethylase 5C |
| SMC1A | Xp11.22 |  | 0.625 |  | 0.575 | Structural Maintenance of Chromosomes 1A |
| KIF4A | Xq13.1 |  | 0.333 |  | 0.080 | Kinesin Family Member 4A |
| TEX11 | Xq13.1 |  | 0.5 |  | 0.541 | Testis Expressed Sequence 11 |
| XIST | Xq13.2 | ncRNA | 1 |  | 0.998 | X Inactive (Non-Protein Coding) |
| JPX | Xq13.2 | ncRNA | 0.667 |  | 0.533 | Nonprotein-coding RNA |
| FTX | Xq13.2 | ncRNA | 0.167 |  | 0.072 | XIST Regulator (Non-Protein Coding) |
| BRWD3 | Xq21.1 |  | 0.5 |  | 0.667 | Bromodomain & WD Repeat Domain Containing 3 |
| APOOL | Xq21.1 |  | 0.167 |  | 0.014 | Apolipoprotein O-Like |
| DACH2 | Xq21.2 |  | 0.333 |  | 0.295 | Dachshund Family Transcription Factor 2 |
| DIAPH2 | Xq21.33 |  | 0.143 |  | 0.076 | Diaphanous-Related Formin 2 |
| ZCCHC16 | Xq23 |  | 0.5 |  | 0.632 | Zinc Finger, CCHC Domain Containing 16 |
| XIAP | Xq25 |  | 0.182 |  | 0.366 | X-Linked Inhibitor of Apoptosis, E3 Ub Ligase |
| SLC9A6 | Xq26.3 |  | - | S | 1 | Solute Carrier Family 9 Member A6 |

<sup>a</sup>Escaper Score according to the strict protocol (DPs) for 25 single lymphoblast cells. The score (0-1.0) is based on scoring genes by data points (DPs) with a paternal allele presence out of total DPs. <sup>b</sup>Genes with a single (S) data point (DP) support. The rest of the genes are supported by multiple DPs according to the strict protocol. <sup>c</sup>Escaper Score according to the relaxed (reads' count) protocol. The ratio (0-1.0) is based on the fraction of the paternal allele in view of the total sum of reads per gene.

### Expansion the list of escapee by read counts

The discrete nature of DP labels for informative SNPs allows analyzing each cell as an independent data source and infer properties of cell variability and consistency. However, this analysis ignores the actual level of expression and the statistical power of some DPs. For example, SLC25A6 is supported by 7880 reads (Supplemental Table S7) that are associated with only two informative SNPs. We reanalyzed the data by adopting a relaxed protocol based on read counts of all informative SNPs per gene (see Materials and Methods). This relaxed protocol (with a minimal threshold of >=7 reads from a paternal allele for a gene) retrieved 31 escapees, which include all the 25 escapees obtained by the strict protocol (Table 2). Applying the same thresholds for analyzing the escapees from the pool data (Pool30 and Pool100) resulted in a small subset of the escapee genes identified by the single cell unified analysis ((7-8 genes by the relaxed protocol, Supplemental Table S8). We expect an improvement in the discovery rate by increasing the number of analyzed single cells, and by having a denser map of informative SNPs.

The higher sensitivity of the relaxed protocol relative to the strict one allowed to separate the apparently monoallelic expression that may result from expression bursts (Islam *et al.* 2011; Borel *et al.* 2015) or from a genuine phenomenon of XCI. We compared reads’ counting for Chr17 for single cells (Figure 5A, Supplemental Table S7) and for Pool30 (Figure 5B, Supplemental Table S8). A simple linear regression for gene expression from the two alleles shows a perfect fit line of 0.995 and a correlation confidence of R^2^ = 0.718. As expected, the correlation confidence for the Pool30 data reached almost a perfect correlation (R^2^ = 0.909, Figure 5B). Same trend was associated with Pool100 analysis (Supplemental Table S8). However, the regression line for ChrX has a much lower fit (y=0.29) with a correlation coefficient of R^2^ = 0.23 (Figure 5C). The slope is indicative of the bias toward maternal expression (x-axis, Figure 5C). Importantly, the correlation coefficient of Pool30 data remained poor (R^2^ = 0.32, Figures 5D). These results confirm that the read counting from ChrX is fully explained by XCI phenomenon and validate the identification of escapee genes (Figures 5C-5D).

**Figure 5.**
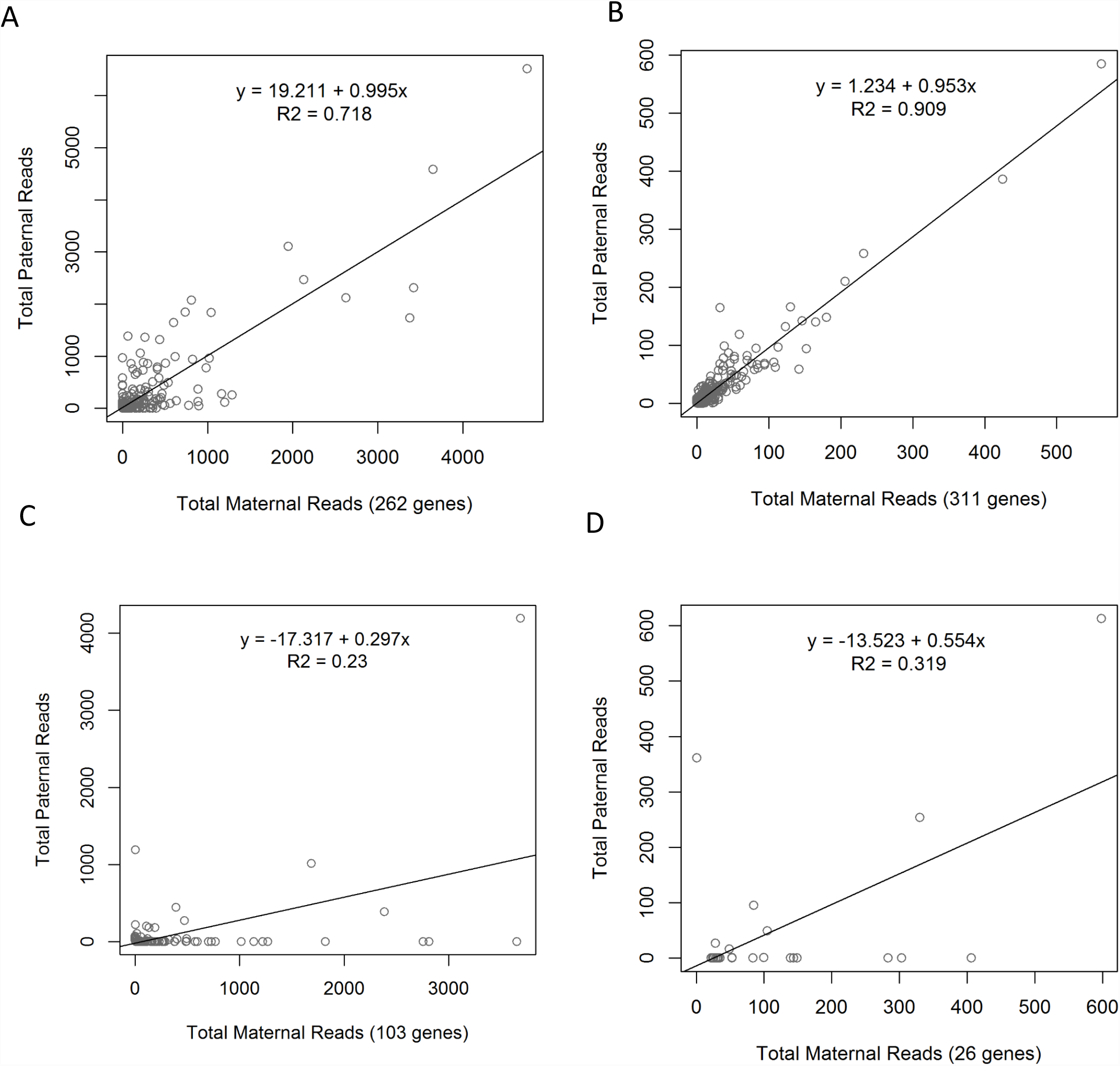
Correlation between the expression from paternal and maternal allelles. The scatter plots show the expression levels of genes by the number of reads associated with the maternal (x-axis) and paternal (y-axis) haplotypes. Only genes supported with >=7 reads are listed. The number of genes for each scatter plot is indicated (on the x-axis, in parenthesis). Data shown are from Chr17 based on single cells (A) and pool30 (B) analysis. Data shown are from ChrX based on single cells (C) and pool30 (D). Note that the number of reads for the pool data is 5-10 folds smaller with respect to the data extracted from the single cells (A, C).

### Only few genes are exclusive escapees - a single cell view

Current estimates suggest about 20% of ChrX human genes to be escapees. This estimate is according to a literature-based catalog that synthesized several reliable publications on escapees and inhibited genes. The indirect methods that were considered in identifying escapees include human-mouse cell hybrids, SNP array, epigenomic marks and expression biases between genders (Balaton *et al.* 2015). We tested the match between identified escapees and inhibited genes from our single cell analysis in view of current knowledge of the annotated genes from ChrX (total 1144 genes) (Balaton *et al.* 2015). The detailed annotation scheme (with 9 different annotations, see Materials and Methods) marked 17% of the genes with an escapee phenomenon (168/630 annotated genes), with only 4.5% of the genes are labelled as exclusive escapees. Based on these annotations, we calculate the statistical significance of the overlap between escapees identified in our study and the unified literature-based catalog (Table 3). We found a statistical significant correspondence between our single-cell based lists (Table 2) and the literature-unified catalog. The calculated P-values range from 1.45E-05 to 1.76E-7 for the strict and relaxed protocols, respectively. PAR genes were identified among the identified escapees (Table 2). Therefore, we critically tested the possibility that the strong statistically significance rely entirely on successful identification of PAR genes. We repeated the statistical test following removal of the PAR genes. Still, the significance of the analysis remains high (P-values are 1.93E-03 and 4.7E-06, for the strict and relaxed protocols, respectively). We conclude that the list of escapees obtained from single cells analysis agree with current knowledge on escapees (See Table 3). Noticeably, by increasing the threshold for the relaxed protocol from >7 to >14 parental reads per gene the specificity of escapee identification was increased with 20/26 identified genes matched the escapee-related annotation (P-value 5.94E-09). *XIST*, the ncRNA that drives ChrX silencing was counted among the genes that did not matches the annotated escapee catalog. Actually, data from single cells clearly show that *XIST* matches a characteristic of escapee, as its expression is exclusively from the Xi chromosome (i.e., parental in the case of the lymphoblasts). Two additional genes (*TMSB4X* and *TEX11*) that were identified as escapees (Table 2) lack information in the literature, and thus were excluded from the statistical analysis. The confident measurement for these genes strongly support their identity as genuine escapee genes.

**Table 3.** Statistical significance by the hypergeometric distribution for the intersection of literature based escapee catalog and escapee lists derived from this study

| Protocol | N | k | n | x | Comment [n] | P-value ($\geq x$) |
| --- | --- | --- | --- | --- | --- | --- |
| Strict | 630 | 168 | 23 | 15 | by SNP DP $\geq 2$ | 1.447E-05 |
| Strict (no PAR) | 608 | 146 | 17 | 9 | by SNP DP $\geq 2$ | 1.935E-03 |
| Relaxed | 630 | 168 | 29 | 20 | by reads, paternal reads $> 7$ | 1.757E-07 |
| Relaxed | 630 | 168 | 26 | 20 | by reads, paternal reads $> 14$ | 5.936E-09 |
| Relaxed (no PAR) | 608 | 146 | 19 | 13 | by reads, paternal reads $> 14$ | 4.717E-06 |

N, k, n and x refer to standard hypergeometric notations (see Materials and Methods). N includes the summary table based on several datasets and unified annotations according to (BALATON AND BROWN 2016). The catalog also contains 22 PAR annotated genes.

What can we learn about escapers’ properties with regard to their expression pattern? It was proposed that for validated escapees, the expression from the Xi is strongly suppressed with respect to the expression from Xa. We thus tested the fraction of the paternal expression of identified escapees from the lymphoblasts (Figure 6A). We observed that genes showing mostly paternal reads are in general low expressing (Supplemental Table S7). This is in agreement with the observation that associates the lower expressing allele to the inhibited chromosome (Carrel and Willard 2005; Zhang *et al.* 2013). An interesting case is that of the *XIST,* which is characterized by an extremely high paternal (Xi) expression, as anticipated from its role (Plath *et al.* 2002; Agrelo *et al.* 2009). It is likely that some of the low expressing genes that show purely maternal expression may still be false negatives.

**Figure 6.:**
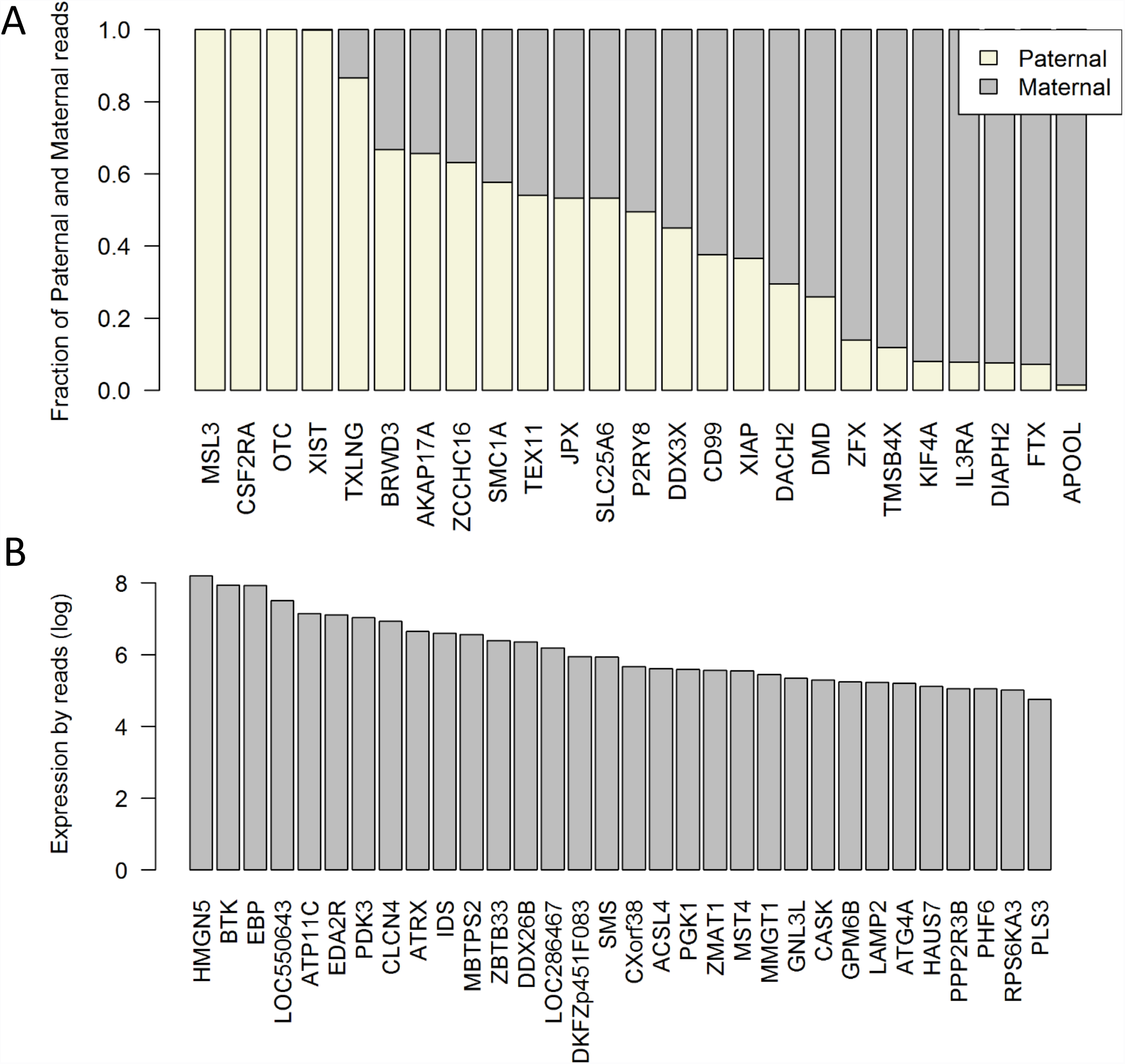
Expression levels of escapees and inhibited genes analyzed from single cell lymphoblasts. (A) Escapees are partition into labels according to the haplotype source of the reads as paternal (beige) and maternal (gray) reads. (B) The total reads assigned to 32 inactivated genes in which each gene is expressed by at least 100 reads from the maternal Xa with minimal evidence for paternal reads (< 5 reads). A log scale indicates the expression level.

We anticipate that varying expression levels of the candidate escapees (Supplemental Table S5) may explain variations in phenotypes and clinical outcomes in women and men with an altered appearance of sex chromosomes. In this study we have not discussed inhibited genes and focused on escapee identification. However, we were able to determine high confident inhibited genes by setting a high threshold of >100 maternal reads. Figure 6B shows the expression level of these inactivated genes. We report on 32 inhibited genes with high probability, obviously the actual number of inhibited genes is much larger.

Careful analysis of the identified escapees (25 of high confidence, Table 2) suggests that the majority of them have a mixed tendency to act as escapees and inactivated genes or identified with conflicting identity (Supplemental Table S7). This finding is in accord with the emerging notion that escaping X-inactivation is a condition dependent property (e.g., by tissues and human populations), supporting the non-deterministic nature of escapee genes (Peeters *et al.* 2014)). Exclusive escapees that we identified in the strict analysis and were corroborated by the fibroblasts analysis (Table 1) include the *ZFX*, S*MC1A*, and *DDX3X*. These genes function in binding and regulating of nucleic acids. *ZFX*, *SMC1A*, and *DDX3X* belong to the short list of exclusive escapees (Balaton *et al.* 2015). *ZFX* is a transcriptional regulator and was repeatedly identified as escapee with its homologous gene (*ZFY*) on the Y chromosome. *SMC1A* is part of the cohesion complex that aligns the sister chromatids for correct segregation of chromosomes during division. *DDX3X*, an RNA helicase that function in transcription, splicing and RNA transport and its mutated version leads to mental retardation. In addition to *XIST*, we identified *JPX*, a ncRNA that acts in coordination with *XIST* for ChrX silencing. In summary, among the exclusive escapees that we have identified we noted an abundance of nucleic acid regulators that affect developmental processes.

We illustrated that ChrX genes properties (as escapees or inhibited) are captured at a single cell level, while the sensitivity is drastically reduced by data from pools of cells, including for pools from clonal cells. The list of identified escapees (31 genes, Table 2) and additional identification from fibroblasts (13 genes, Table 1) are mostly located in the p-arm of ChrX. This is in agreement with the observed distribution of escapees along ChrX. The enrichment of escapees in the p-arm reflects the recent evolutionary history of human sex chromosomes. We show that single-cell analysis from RNA-Seq is valuable as a sensitive and robust method for identifying X-inactivation and genes escaping it.

## ACKNOWLEDGEMENTS

We thank Nati Linial and Nadav Brandes for useful discussion and critical comments. We thank Nadav Brandes for critical reading of the manuscript. We thank Matan Avraham for help with the Blast script. We thank Shachar Shohat for help with the SNP calling. We thank Yuval Nevo and the CSE system for technical assistance.

## Competing interest

The authors declare that they have no competing interests.

## Funding

The research was partially supported by the EU-H2020 Elixir-Excelerate.

## Authors’ contribution

KWK and ML wrote the manuscript, performed the design and the analysis. Both authors read and approved the final manuscript.

## Supplementary Figures

**Figure S1.**
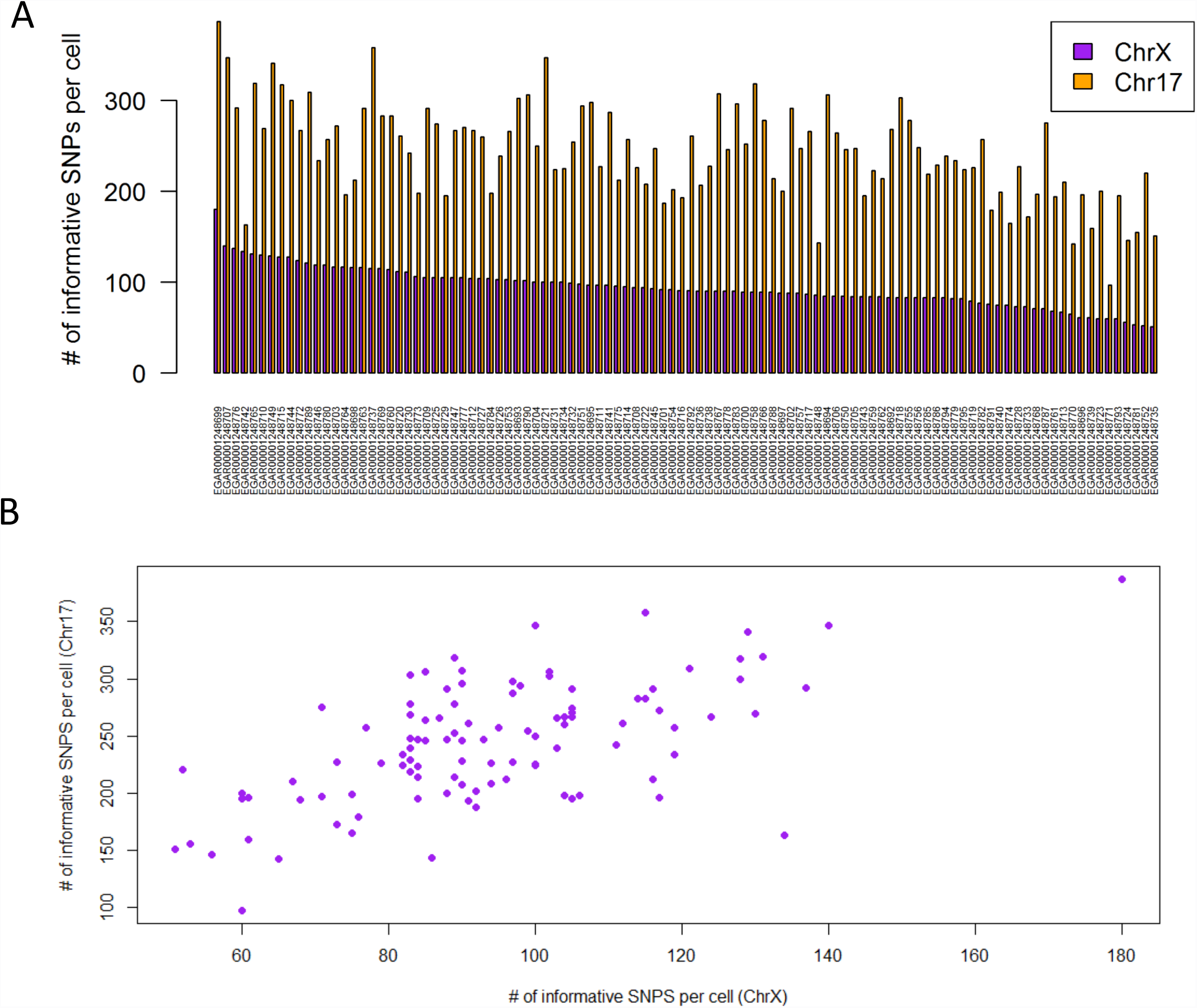
Informative SNPs on both chromosomes of fibroblasts **(A)** The number of informative SNPs for each of the 104 cells from ChrX and Chr17. Data was collected from 104 fibroblast cells of female origin (UCF_1014). The cells’ identifiers are listed in Supplementary Table S1. **(B)** Correlation between the numbers of SNPs for the two chromosomes according to an individual cell. Pearson's correlation of the SNPs for these two chromosomes for all tested cells is r = 0.618, p-value = 2.781e-12.

**Figure S2.**
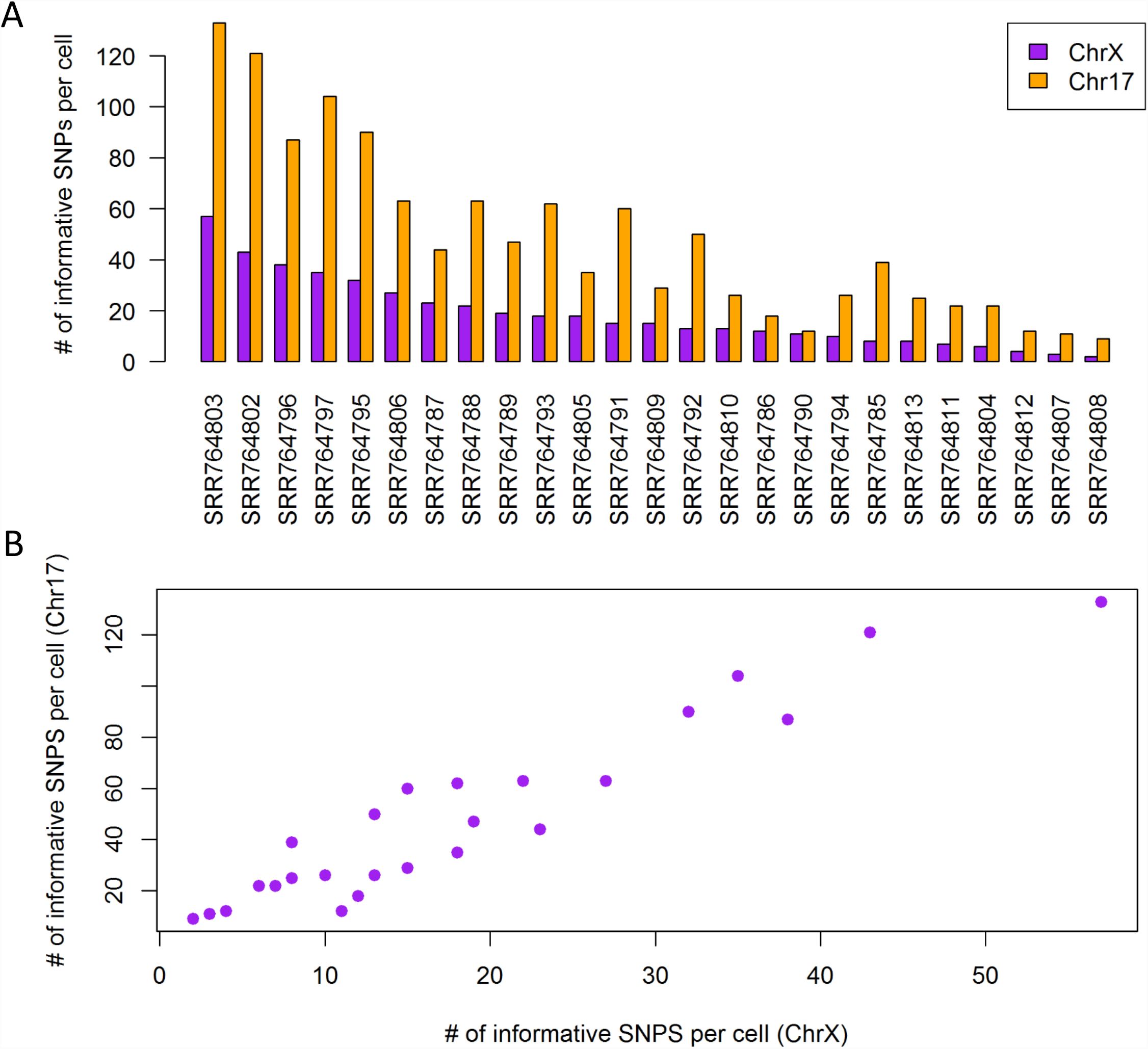
Informative SNPs on both chromosomes of lymphoblasts (A) The number of informative SNPs for each of the 25 cells from ChrX and Chr17. Data was collected from 25 cells of female origin (GM12878 lymphoid cell-line). The cells’ identifiers are listed in Supplementary Table S4. (B) Correlation between the numbers of SNPs for the two chromosomes according to an individual cell. Pearson's correlation of the SNPs for these two chromosomes for all tested cells is r = 0.948, p-value = 5.989e-13.

## Supplementary Tables legends

**Table S1.** Names of RNA-seq names for 104 primary fibroblast UCF_1014 cells. Number of aligned reads and the minimal read limit are indicated.

**Table S2.** List of informative SNPs assigned to ChrX and Chr17 from 104 primary fibroblast single cells. The legends for the columns are: contig-chromosome; position-position on the chromosome; variantID – the official ID of the SNP; refAllele – the reference allele; altAllele - the alternative allele; Genes – the genes corresponding to the SNPs (empty cells indicate intergenic positions). For the other columns, each column represents a single cell. The rows indicate the different informative SNPs. For the labelling of the SNPs we used the following color-code: Gray-no reads; Purple-Reference allele (#Alt reads/(#Ref + #Alt reads) <=0.25); light purple-biallelic dominated by the Reference allele (0.25<#Alt reads/(#Ref + #Alt reads) <=0.5); Brown-balanced expression leaning towards Alternative (0.5<#Alt reads/(#Ref + #Alt reads) <=0.75); Yellow-Alternative (#Alt reads/(#Ref + #Alt reads)>0.75).

**Table S3.** Gene-centric view on Allelic Ratio for primany fibroblasts. Columns correspond to: Genes – genes containing the SNPs; readsRef – reads assigned to the Reference allele; readsAlt – reads assigned to the alternative allele; ReadsTotal – the total number of reads aligned to this gene; AltRatio -the ratio of alternative reads form total reads; RefDP – number of DP determined as the Reference allele; AltDP -number of DP determined as the Alternative allele; BalancedDP-number of DP determined as biallelic; TotalDP-total number of DPs in gene; BalancedRatio – ratio of balanced DPs out of total DPs; Strict_Protocol_Identification -the identification of the gene as in ChrX inactivated or Escaper, or in Chr17 as biallelic or monoallelic. Gene supported by a single DB is marked as ‘less than 2 DP’.

**Table S4.** Names and database indexes of RNA-seq datasets for the 25 single GM12878 lymphoid cells and the pool samples. The number of reads for raw data, filtered by FASTAX, aligned to ChrX and Chr17, and aligned to only one location are also shown.

**Table S5.** List of informative SNPs assigned to ChrX and Chr17 from 25 single cells. Each chromosome is in a different sheet. First 6 columns correspond to: chr – chromosome; snpRef – location of SNP on Reference genome; paternal – location of SNP on paternal genome; maternal – location of SNP on maternal genome; names - names of SNPs in our GTF file; GenesOfSnps – the gene that contains each SNP. For the other columns, each column represents a single cell. For the labelling of the SNPs we used the following color-code: gray-no reads; purple-maternal (#paternal reads/#maternal reads <=0.25); light purple-mostly maternal (0.25=#paternal reads/#maternal reads<=0.5); brown-balanced expression leaning towards paternal (0.5<#paternal reads/#maternal reads=<0.75); yellow-paternal (#paternal reads/#maternal reads>0.75).

**Table S6.** List of informative SNPs assigned to ChrX and Chr17 on pool30 and pool100. Each chromosome in each pool analysis (Pool30 or Pool100) is in a different sheet. The other settings are as in Table S5.

**Table S7.** Genic centred allelic determination of Allelic Ratio of lymphoblast 25 single cells. Each chromosome is in a different sheet. Columns correspond to: Genes – Genes containing the Snps; readsPat – reads assigned to the Paternal allele; readsMat – reads assigned to the maternal allele; ReadsTotal – the total number of reads aligned to this gene; PatRatio – The ratio of paternal reads out of the total; Relaxed_Protocol_Identification – Relaxed protocol identification of genes for ChrX as Escaper or inhibited and in Chr17 as Paternal Maternal or bi-allelic expressed; MaternalDP – number of DP determined as Maternal allele; PaternalDP – number of DP determined as Paternal allele; BalancedDP -number of DP determined as bi-allelic; TotalDP - total number of DPs in gene; EscaperRatio – ratio of paternal and balanced DPs out of total DPs (The DP score); Strict_Protocol_Identification – The identification of the gene as in ChrX inactivated or Escaper, or in Chr17 as bi-allelic, maternal or paternal allele (If less then 2 DPs were indicative ‘less than 2 DP’ is indicated); protocols_Used – protocols used to reach final conclusion where R stands for Relaxed and S for Strict; overall_Identification – identification and by how many methods it was identified.

**Table S8.** Genic centred allelic determination of Allelic Ratio of lymphoblast Pool30 and Pool100. Each chromosome in each pool analysis (Pool30 or Pool100) is in a different sheet. The other settings are as in Table S7.

